# Applying particle filtering in both aggregated and age-structured population compartmental models of pre-vaccination measles

**DOI:** 10.1101/340661

**Authors:** Xiaoyan Li, Alexander Doroshenko, Nathaniel D. Osgood

## Abstract

Measles is a highly transmissible disease and is one of the leading causes of death among young children under 5 globally. While the use of ongoing surveillance data and – recently – dynamic models offer insight on measles dynamics, both suffer notable shortcomings when applied to measles outbreak prediction. In this paper, we apply the Sequential Monte Carlo approach of particle filtering, incorporating reported measles incidence for Saskatchewan during the pre-vaccination era, using an adaptation of a previously contributed measles compartmental model. To secure further insight, we also perform particle filtering on an age structured adaptation of the model in which the population is divided into two interacting age groups – children and adults. The results indicate that, when used with a suitable dynamic model, particle filtering can offer high predictive capacity for measles dynamics and outbreak occurrence in a low vaccination context. We have investigated five particle filtering models in this project. Based on the most competitive model as evaluated by predictive accuracy, we have performed prediction and outbreak classification analysis. The prediction results demonstrated that this model could predict the measles transmission patterns and classify whether there will be an outbreak or not in the next month (Area under the ROC Curve of 0.89). We conclude that anticipating the outbreak dynamics of measles in low vaccination regions by applying particle filtering with simple measles transmission models, and incorporating time series of reported case counts, is a valuable technique to assist public health authorities in estimating risk and magnitude of measles outbreaks. Such approach offer particularly strong value proposition for other pathogens with little-known dynamics, critical latent drivers, and in the context of the growing number of high-velocity electronic data sources. Strong additional benefits are also likely to be realized from extending the application of this technique to highly vaccinated populations.

**Author summary:** Measles is a highly infectious disease and is one of the leading causes of death among young children globally. In 2016, close to 90,000 people died from measles. Measles can cause outbreaks particularly in people who did not receive protective vaccine. Understanding how measles outbreaks unfold can help public health agencies to design intervention strategies to prevent and control this potentially deadly infection. Although traditional methods – including the use of ongoing monitoring of infectious diseases trends by public health agencies and simulation of such trends using scientific technique of mathematical modeling – offer insight on measles dynamics, both have shortcomings when applied to our ability to predict measles outbreaks. We seek to enhance the accuracy with which we can understand the current measles disease burden as well as number of individuals who may develop measles because of lack of protection and predict future measles trends. We do this by applying a machine learning technique that combines the best features of insights from ongoing observations and mathematical models while minimizing important weaknesses of each. Our results indicate that, coupled with a suitable mathematical model, this technique can predict future measles trends and measles outbreaks in areas with low vaccination coverage.

## Introduction

Measles is a highly contagious viral disease. It remains one of the leading causes of death among young children globally, and has imposed a significant morbidity and mortality burden where vaccination coverage is low [1]. About 30% of all cases of measles have one or more complications including diarrhea, otitis media, pneumonia or encephalitis. Measles mortality was estimated to be 0.2% in the United States between 1985 and 1992 [2]. Prior to widespread vaccination, measles caused an estimated 2.6 million deaths each year [1]. In 2016, approximately 89,780 people died from measles – mostly children under the age of 5 [1]. The Americas has become the first region in the world to have eliminated measles [3], however importation of cases from other regions leads to outbreaks in unimmunized and under-immunized populations. Understanding measles epidemic patterns can aid in forecasting and help public health agencies to design intervention strategies to prevent and control it, such as setting outbreak response measures, setting vaccination targets, and allocating financial and human resources, etc.

Traditionally, many public health agencies seek to anticipate future transmission of infectious diseases based upon time series of reported incident case counts, derived from public health surveillance. In recent years, simulation models [4, 5] have increasingly been used to predict the spread of measles within the population, and the potential impact of interventions on that spread. Although both these methods have their benefits, and have played an important role in estimating and predicting measles epidemic pattern, each suffers from notable limitations.

In many jurisdictions worldwide, measles is a notifiable illness under respective Public Health Acts, and surveillance reports are collected and examined on a regular basis. These data – often summarized as epi-curves – have offered great value for public health agencies seeking to understand trends in measles incidence and anticipate future evolution. For example, identifying and confirming suspected measles cases through surveillance allows early detection of outbreaks, and supports comparing recent activity to reference periods. However, the reported time series data are often quite noisy, particularly for contexts marked by smaller population or low incidence or diagnosis rates. Such reports are also delayed, and may be incomplete, since families of some of those infected elect not to seek care. When used unassisted, time series lack the capacity to provide quantitatively rigorous insight into the future evolution of the observed patterns -– evolution that will be shaped by a complex combination of local circumstances, including birth rates, contact patterns, and the size of latent factors such as the local pool of susceptibles as affected by vaccination rates, immunity acquired after contracting disease and recent history of outbreaks. Perhaps most importantly when considered as a tool for planning, use of time series alone cannot itself be used to investigate counterfactuals, such as how an outbreak response immunization campaign, enhanced contact tracing, advisories, exclusions from schools, day care and workplaces or social distancing measures are likely to affect future incidence dynamics.

Dynamic modeling has played a significant role in providing insight to the measles outbreak dynamics [4,6–15]. Most such contributions employing models seek to incorporate some aspects of local epidemiology, <and often draw on surveillance data. While powerful tool for investigating counterfactuals, such models also suffer from an essential set of shortcomings. Firstly, while dynamic models are commonly calibrated to empirical data, this process is typically undertaken on a one-time basis, and with significant human involvement. It is rare for a dynamic model to incorporate ongoing arriving ground data; and while systems doing so can be found for other infectious diseases [40], the authors are not aware of any such support for dynamic modeling of measles. More profoundly, a dynamic model of necessity represents a simplified characterization of processes in the real world. Inevitably, such models often omit, simplify and misestimate some factors. These drawbacks and the infeasibility of anticipating the realized outcome of factors represented as stochastic in the model will inevitably lead the model to diverge from ground truth data. While calibration can allow for estimation of model parameters, it provides weak support for ongoing estimation of latent state needed to keep the state of a model aligned with observations on an ongoing basis.

This paper seeks to support more accurate estimation and prediction of measles dynamics by applying a computational statistics technique that combines the best features of insights from ongoing (although noisy) empirical data and dynamic models (although fraught by systematic errors, omissions, and stochastic divergence over time) while mitigating important weaknesses of each. The use of sequential Monte Carlo methods in the form of particle filtering [16–28] has provided an effective and versatile approach to solving this problem in other infectious diseases, such as the influenza. Specifically, this paper investigates the combination of particle filtering methods with a compartmental model (SEIR model) of measles to recurrently estimate the latent state of the population with respect to the natural history of infection with measles, to anticipate measles evolution and outbreak transitions in pre-vaccination era.

## Results

### Model characterization and discrepancy comparison

To research on the performance of incorporating particle filtering into the compartmental model, we have built 7 models, including 1 normal deterministic model, 1 calibration model and 5 particle filtering models. In the particle filtering models, the number of particles are all 5000. These models respectively listed as follows:

1. *Pure_aggregate_*. It is simply the deterministic SEIR model with aggregate population, see the Eq (1). The value of the initial infectious population is 90, the initial susceptible population is 89910, the initial exposure population is 0, and the initial recovered population is 773545. The values of *β* and reporting rate *C_r_* are 50 and 0.11, respectively.
2. *Calibrated_aggregate_*. It is the calibration model of the SEIR model with aggregate population. However, the relatively uncertain and stochastic parameters, including initial infectious population, initial susceptible population, the parameter *β* and reporting rate *C_r_* are obtained by calibration. Finally, they are relatively 930, 89070, 49.598 and 0.119. The initial exposure population is 0, and the initial recovered population is 773545.
3. *PF_aggregate_*. The particle filtering model with homogeneous mixing of all population.
4. *PF_age_5_monthly_*. The age structure model where the child age group includes those less than 5 years old, and only incorporated with the monthly reported empirical data.
5. *PF_age_5_both_*. The age structure model where the child age group includes those less than 5 years old, and incorporated with both the monthly reported and yearly reported age group empirical data.
6. *PF_age_15_monthly_*. The age structure model where the child age group includes those less than 15 years old, and only incorporated with the monthly reported empirical data.
7. *PF_age_15_both_*. The age structure model where the child age group includes those less than 15 years old, and incorporated with both the monthly reported and yearly reported age group empirical data.

By comparing the discrepancy of these models, we sought to identify the model offering the greatest predictive validity. We then used the most favorable model to perform prediction analysis. To assess model results, each of the five particle filtering models was run 5 times with random seeds generated from the same set. We then calculated the average and 95% confidence intervals of the mean discrepancy.

Table 1 and Fig 1 show the comparison of the discrepancy among the seven models. It is notable that the yearly discrepancy is not available for the aggregated population models. The results demonstrate that the particle filtering models strongly decrease the model discrepancy. This indicates that incorporating particle filtering in the compartmental model of measles could help to improve the simulation accuracy. Secondly, the results suggest that the age structure particle filtering models perform better (as measured by discrepancy) than the aggregated population model, because the monthly discrepancy of all the four stratified age group models are smaller than the aggregated population model. Thirdly, an appropriate splitting of the age groups is also important in improving the simulation results of the models. Table 1 and Fig 1 indicate that the discrepancy of stratified age group models splitting the age group at age 15 years are all smaller than the models splitting the age group at age 5 years. Finally, results suggest that incorporating both the monthly empirical reported cases and yearly empirical data of each age group may be also helpful in improving the simulation accuracy of the models, but further realizations are required to confirm these results. The results suggest that the model *PF_age_15_both_* offers the minimum discrepancy. It is notable that while aggregate population models cannot be compared directly against the other models in terms of total discrepancy, it suffers from the least favorable score in terms of the metric by which comparisons can be made (the monthly discrepancy).

**Fig 1.**
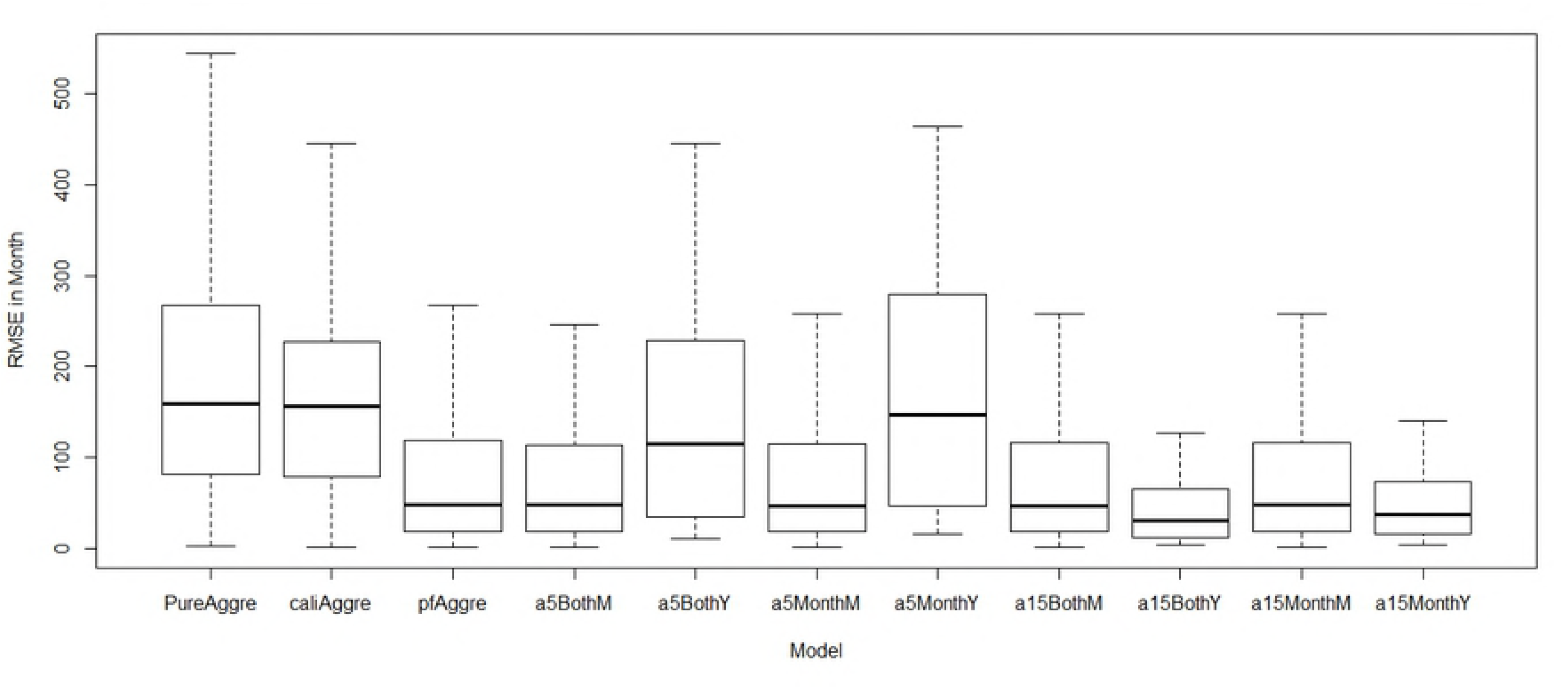
Box plots of monthly and yearly discrepancy of all models by incorporating empirical data across all observation points. Each of the five particle filtering models was run 5 times (the random seed generated from same set). Then the average monthly and yearly discrepancy among these five runs at each time between the particle filtering models and the empirical data were plotted.

**Table 1.** Comparison of the average discrepancy of all seven models by incorporating empirical data across all observation points.

| Model | Monthly | Yearly in Month | Total |
| --- | --- | --- | --- |
| $Pure_{aggregate}$ | 249.0 | NONE | NONE |
| $Calibrated_{aggregate}$ | 207.5 | NONE | NONE |
| $PF_{aggregate}$ | 104.6 (99.4, 109.9) | NONE | NONE |
| $PF_{age\_5\_monthly}$ | 96.1 (91.5, 100.7) | 179.5 (160.6, 198.3) | 275.5 (260.2, 290.9) |
| $PF_{age\_5\_both}$ | 97.8 (94.1, 101.5) | 144.8 (112.5, 177.1) | 242.6 (210.2, 275.1) |
| $PF_{age\_15\_monthly}$ | 95.7 (89.8, 101.6) | 45.9 (39.9, 51.9) | 141.6 (133.8, 149.4) |
| $PF_{age\_15\_both}$ | 96.1 (91.6, 100.5) | 39.9 (34.3, 45.6) | 136.0 (127.6, 144.5) |

Each of the five particle filtering models was run 5 times (the random seed generated from the same set). Shown here are the average and 95% confidence intervals (in parentheses) of the mean discrepancy for each model variant.

### Result Analysis of the minimal discrepancy model

To depict the particle filter results, 2D histograms of reported measles cases calculated by the model and empirical data were plotted. To let the 2D histogram plot characterize the model’s output data with proper weighting in accordance with the principles of importance sampling, we plotted the results of the particles sampled by weights. The resulting plot thus represents an approximation to the probability distribution of reported measles cases characterized by the model. It is notable that the number of particles performed, the number of particles sampled in 2D histograms and the number of particles sampled (also by weight) in calculating the discrepancy in all the models in this paper are all 5000, except where otherwise noted.

Fig 2 and Fig 3 display the prior and posterior results of the particle filtering model *PF_age_ 15_both_* for the entire timeframe, respectively. The results include the 2D histogram of monthly reported measles count and the 2D histogram of yearly reported count of each age group. It is notable that because the weights of particles are updated at each month, the 2D histogram plot giving the prior in Fig 2 is not suitable for the yearly result. Values of empirical data points are shown in red. From them we can see that most of the empirical data are located at the range where the particles exhibit high posterior probability. This reflects the fact that the particles could suitably track the oscillation of the epidemic pattern of measles, given the combination of model prediction and observation-based updating that forms the basis for the particle filter.

**Fig 2.**
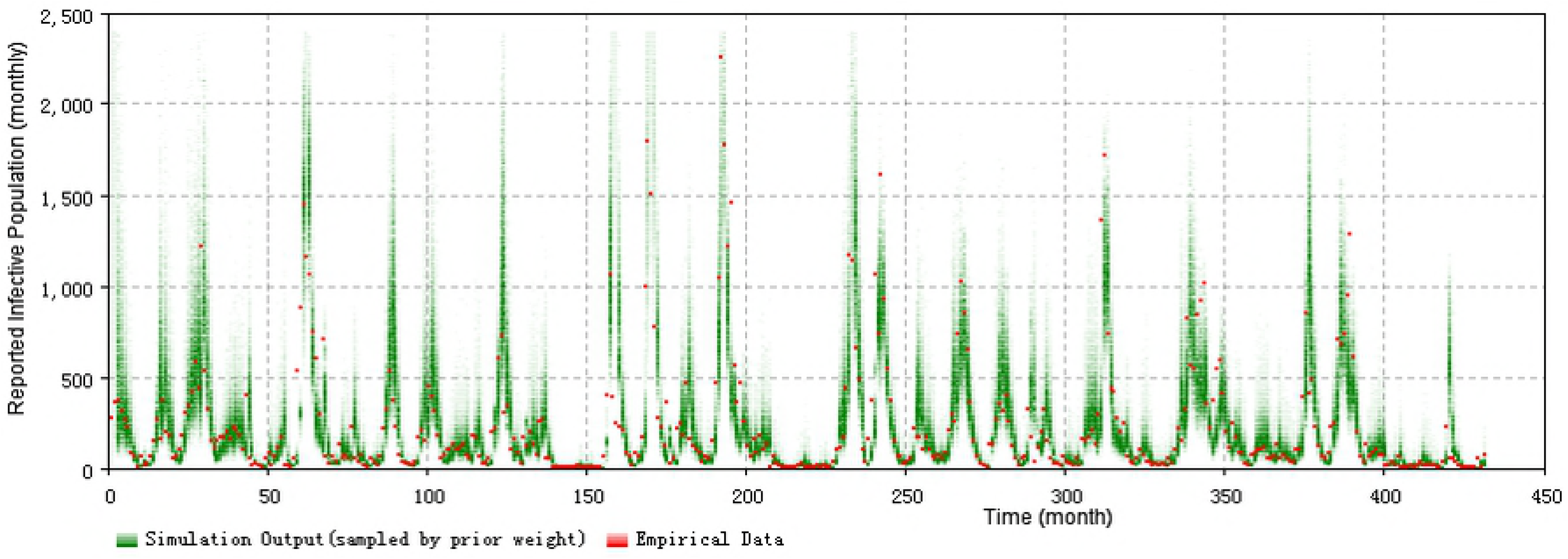
2D histogram prior result of total timeframe of the minimum discrepancy model (monthly).

**Fig 3.**
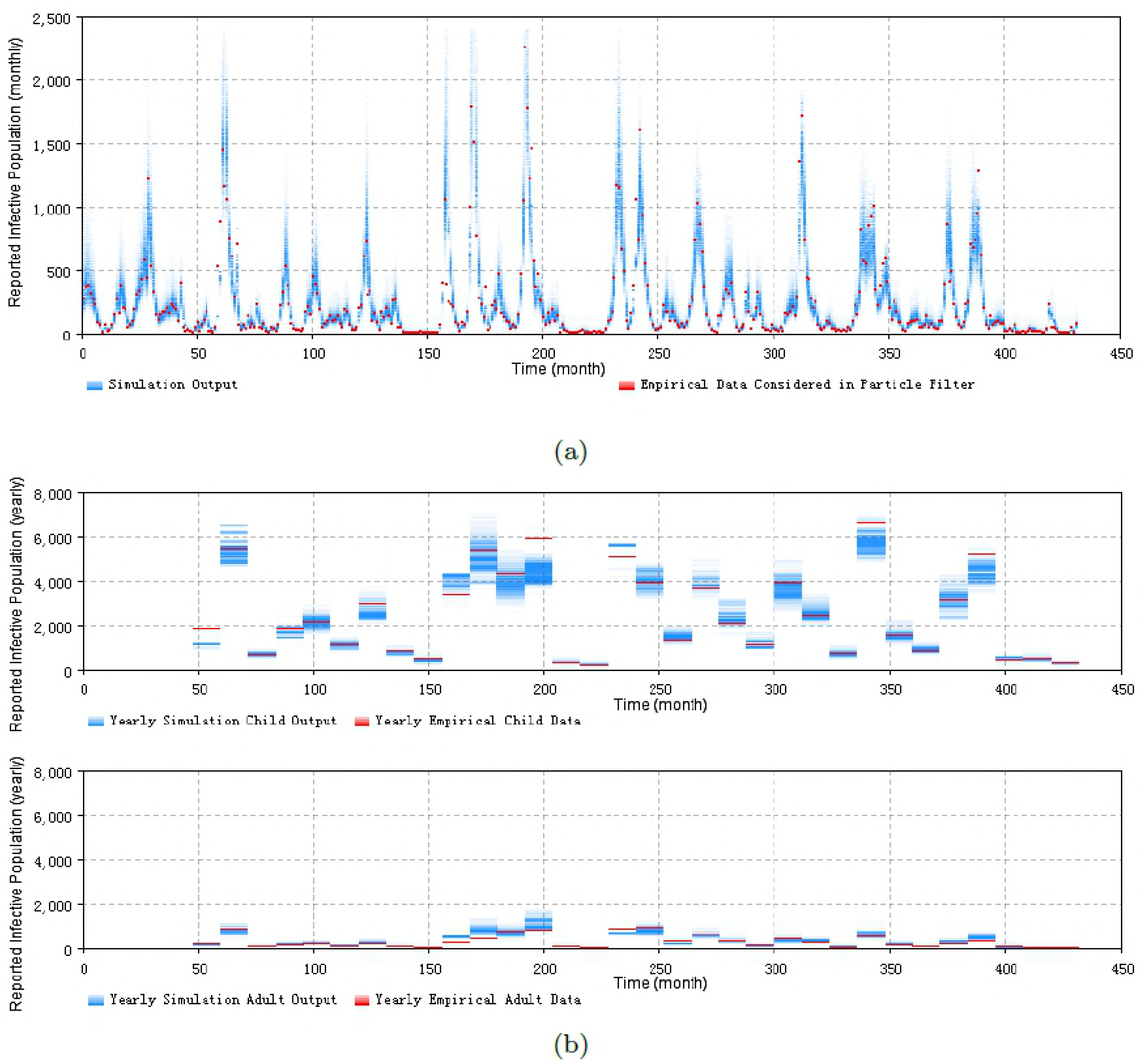
2D histogram posterior result of total timeframe of the minimum discrepancy model. (a) the monthly particle filtering result across all population. (b) the yearly particle filtering result of the children and adult age groups.

It bears emphasis that the results of the particle filtering model sampled in Fig 2 are before the weight update process in each step, while the results in Fig 3 are after the weight update process in each step. The value of sampled particles of Fig 2 spread in a wider range, compared with Fig 3. This difference in dispersion indicates that the weight update process of particle filtering algorithm in this paper has the capability to combine the empirical data to the particle filtering model to constrain the particles in a tighter range as suggested by the empirical data.

Particle filtering models can contribute to the estimation of model states and aid in estimating dynamic model parameters. It is notable that, as is widely the case in dynamic models, the states in the compartmental models are latent (e.g., Susceptible (S), Exposure (E), Infectious (I) and Recovered (R) stocks in the SEIR model Eqs (1)–(5) of measles). What can be empirically observed is the noisy reported measles cases related to the Infectious (I) state. However, the methodology of particle filter provides an approach to estimate (via sampling from) the distribution of values of these latent states. This ability to estimate the value of latent states – such as the reservoir of susceptible people – can aid researchers and public health agencies to in terms of understanding the underlying epidemiological situation from multiple lines of evidence, as constrained by understanding of the structure of the system, as characterized by a dynamic model. To illustrate this, we employ a similar method to the above to plot the 2D histogram of the stocks of susceptible, infectious, exposure and recovered sampled according to importance sampling principles. Fig 4 show the results of the plots. These plots indicate that most of the susceptible, exposure and infectious people are located in the children (less than 15 years) age group, while most of the recovered population are located in the adults (equal and greater than 15 years) age group. This lies in accordance with the expectations for measles transmission in the real world and builds confidence in the capacity of the model to meaningfully estimate latent state. As noted below, estimation of latent state can be an important enabler for understanding of the effects of interventions.

**Fig 4.**
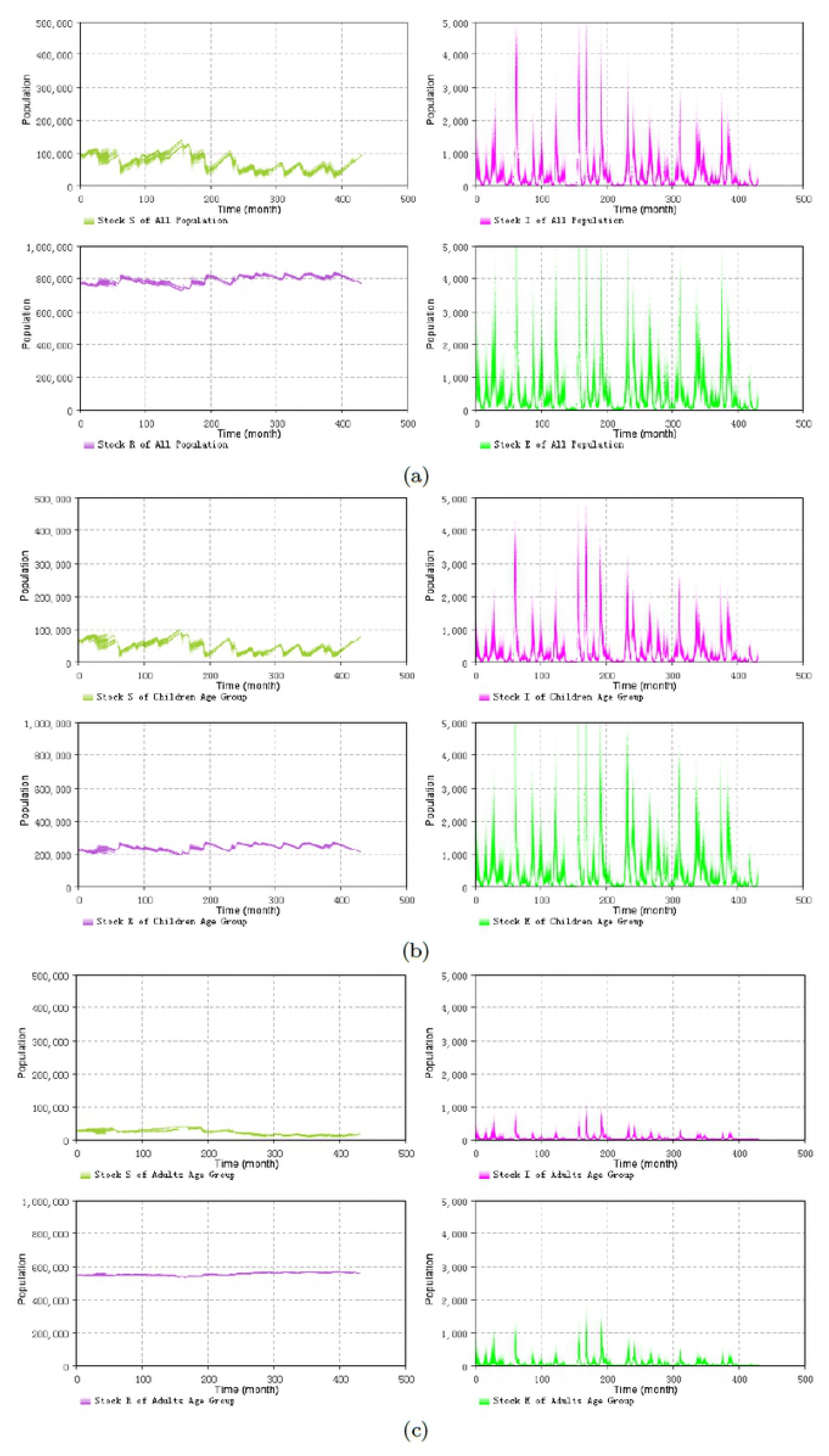
**2D histogram results for the S, E, I, R stocks with different age groups** of the minimum discrepancy model with splitting the age groups at 15 years by incorporating the empirical data across all timeframe. The results shown consider both the yearly and monthly empirical data, with monthly discrepancy 90.7, the sum of all age groups discrepancy in Month is 36.9. (a) across all population. (b) the child age group (those within their first 15 years of life). (c) the adult age group (years 15 and up).

### Prediction results of the minimal discrepancy model

In this section, we assess the predictive capacity of the minimal discrepancy model identified in the previous section. This minimal discrepancy model is still the minimum one among the group of models *PF_age_15_both_*. By changing different Prediction Start Time of T*, we have performed the prediction from different archetypal situations. These situations are listed as follows:

1. Prediction started from the first or second time points of an outbreak.
2. Prediction started before the next outbreak.
3. Prediction started from the peak of an outbreak.
4. Prediction started from the end of an outbreak.

Figs 5–8 display the prediction results of these situations with the monthly 2D histogram of reported cases of the total population. The empirical data having been considered in the particle filtering process are shown in red (incorporated in training the models), while the empirical data having not been considered in the particle filtering process (only displayed to compared with the results of models) are shown in black.

**Fig 5.**
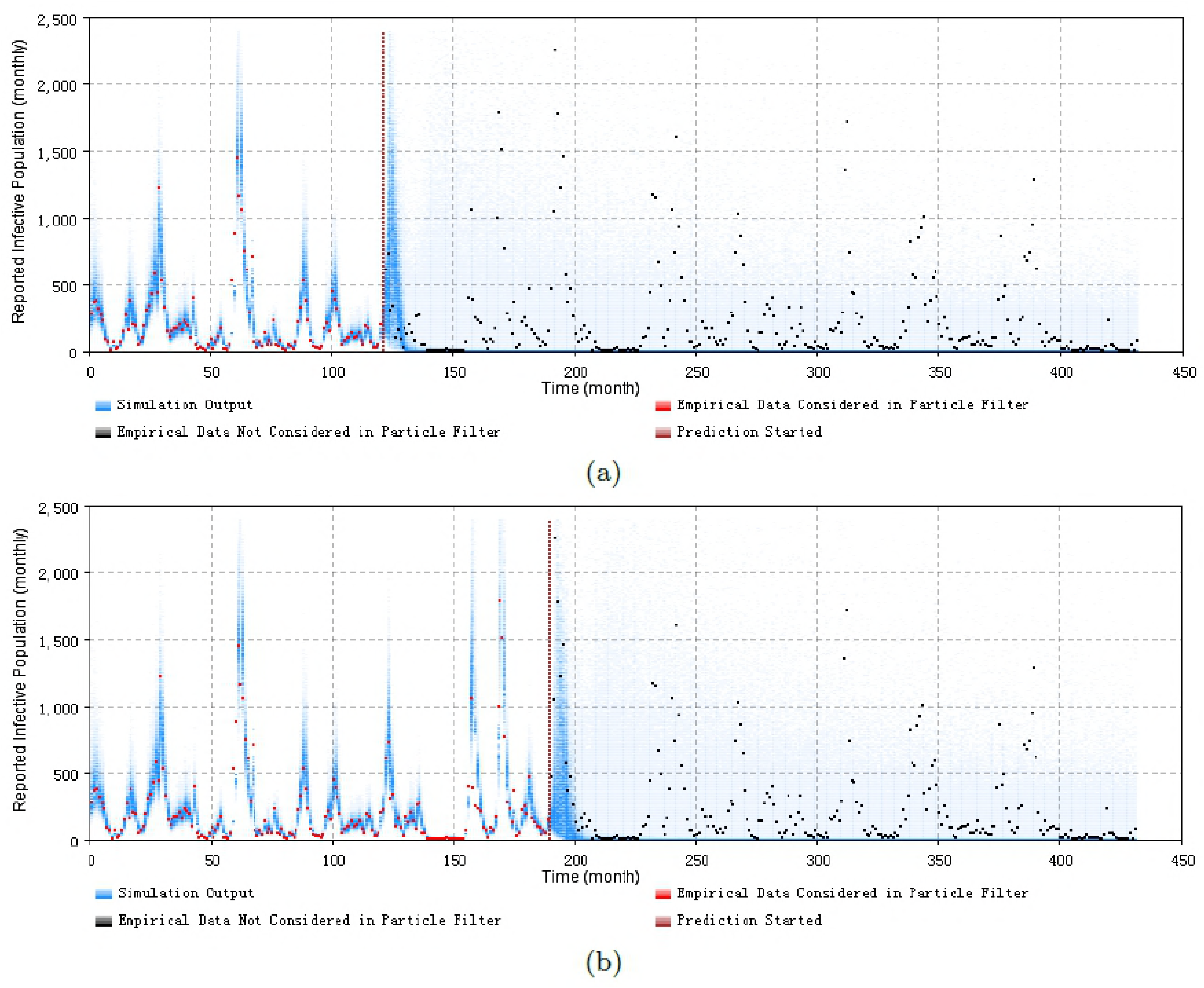
2D histogram of predicting from the first or second time points of an outbreak of the minimum discrepancy model. (a) predicted from the month 121, with monthly prediction discrepancy 306.0, the sum of yearly prediction discrepancy of all age groups per month is 246.7. (b) predicted from the month 190, with monthly prediction discrepancy 320.4, the sum of yearly prediction discrepancy of all age groups per month is 237.2.

**Fig 6.**
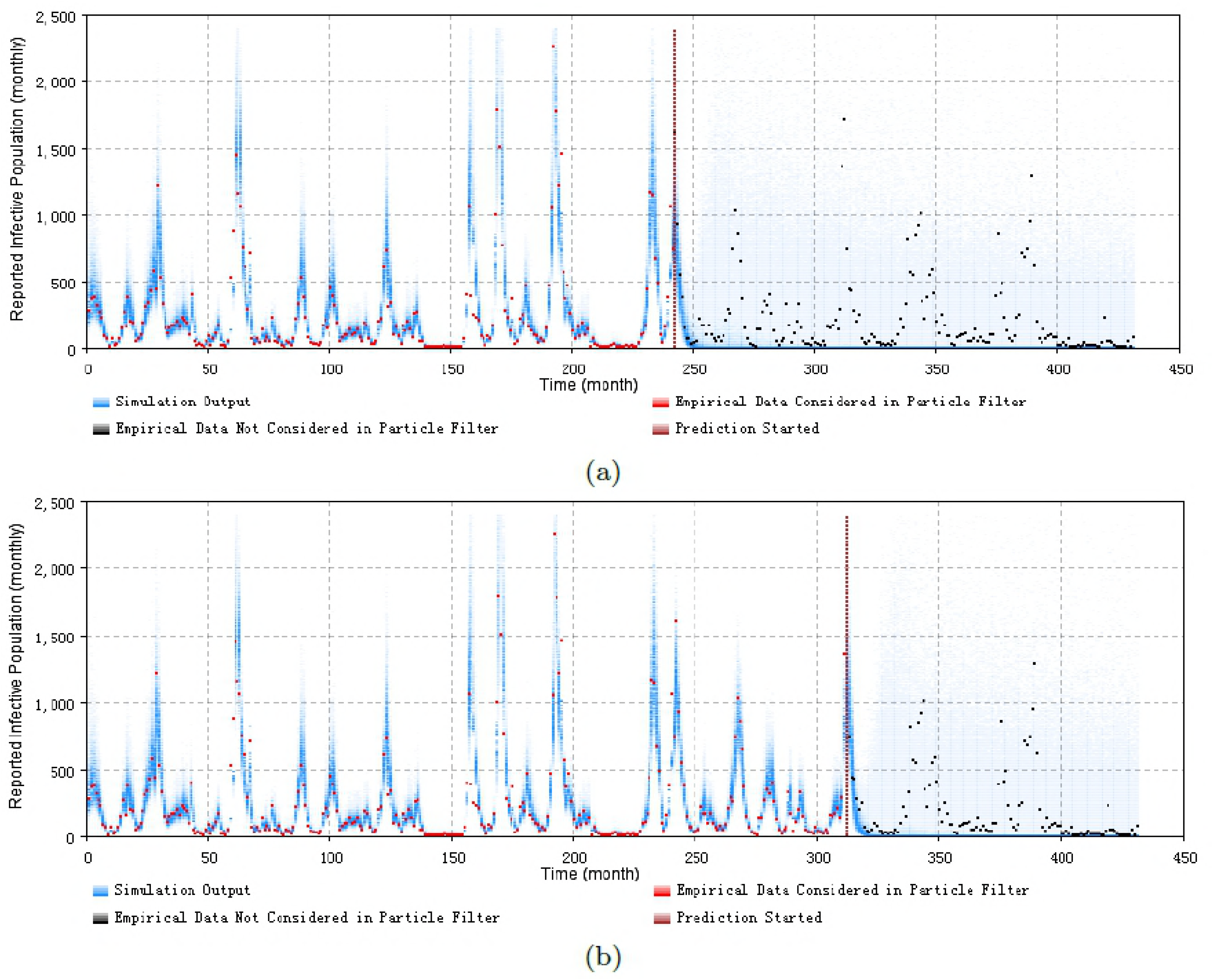
2D histogram of predicting from the peak of an outbreak of the minimum discrepancy model. (a) predicted from the month 242, with monthly prediction discrepancy 305.7, the sum of yearly prediction discrepancy of all age groups per month is 205.2. (b) predicted from the month 312, with monthly prediction discrepancy 306.9, the sum of yearly prediction discrepancy of all age groups per month is 201.6.

**Fig 7.**
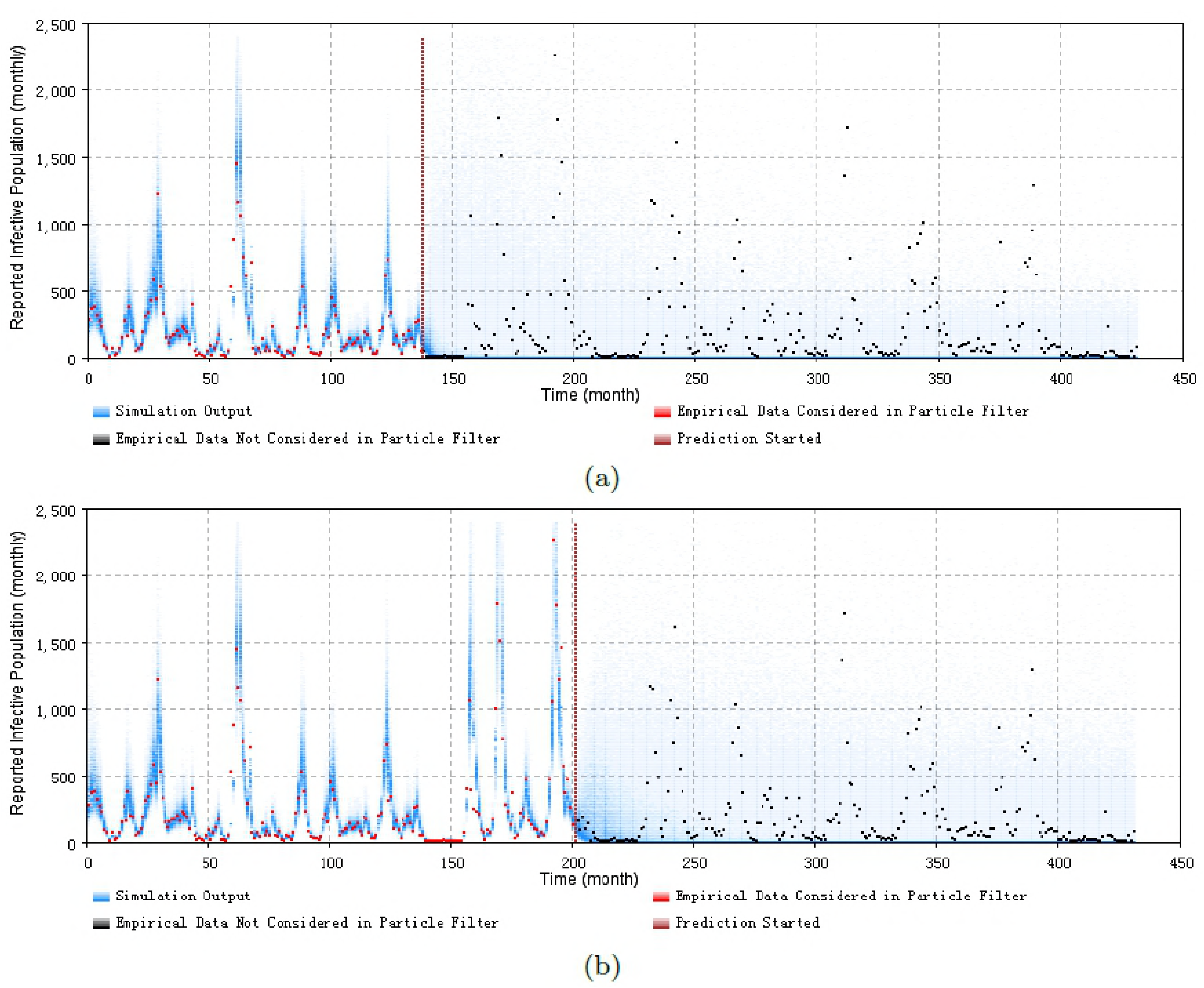
2D histogram of predicting from the end of an outbreak of the minimum discrepancy model. (a) predicted from the month 138, with monthly prediction discrepancy 302.6, the sum of yearly prediction discrepancy of all age groups per month is 248.0. (b) predicted from the month 201, with monthly prediction discrepancy 316.7, the sum of yearly prediction discrepancy of all age groups per month is 217.3.

**Fig 8.**
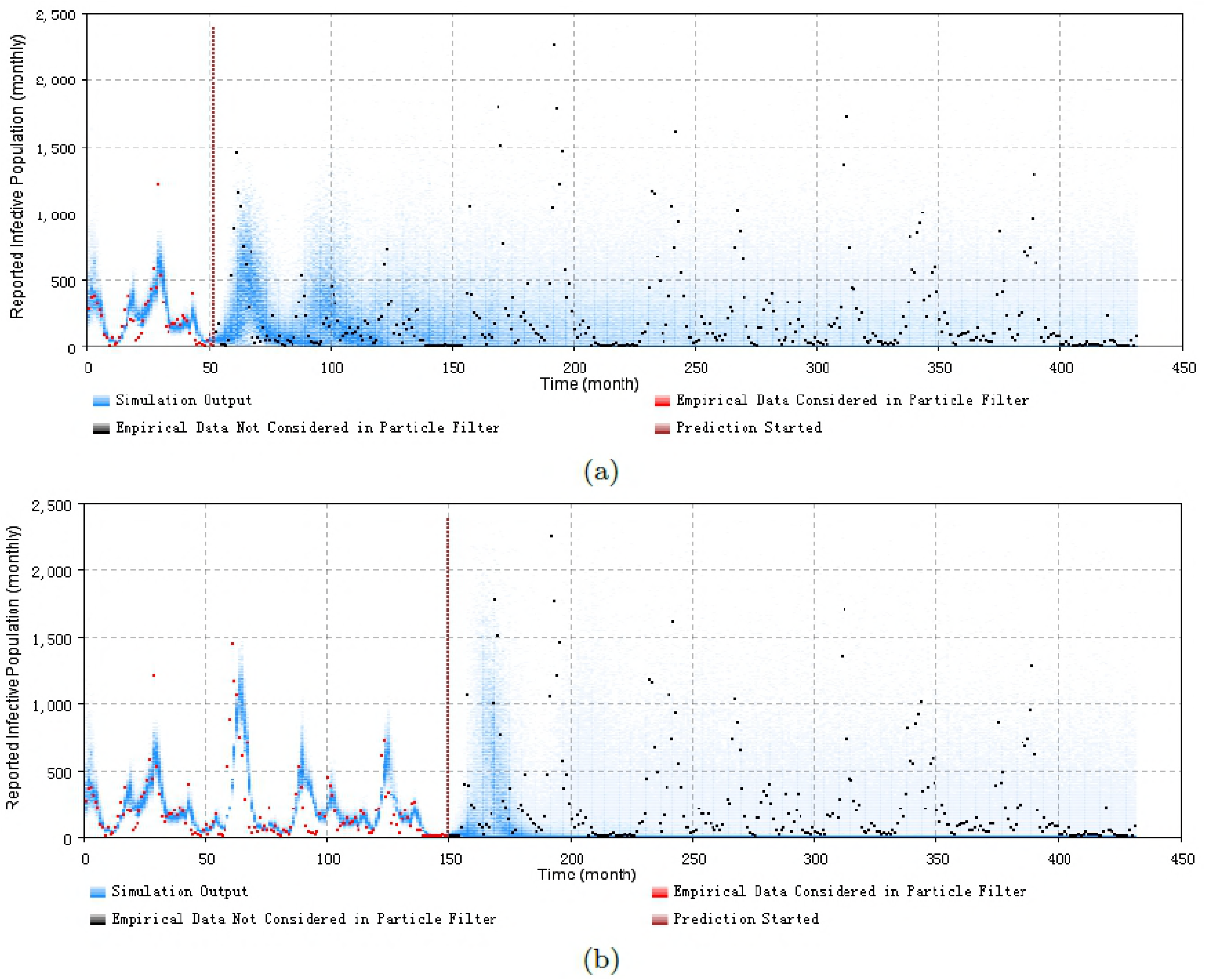
2D histogram of predicting before the next outbreak of the minimum discrepancy model. (a) predicted from the month 52, with monthly prediction discrepancy 324.9, the sum of yearly prediction discrepancy of all age groups per month is 198.8. (b) predicted from the month 150, with monthly prediction discrepancy 353.0, the sum of yearly prediction discrepancy of all age groups per month is 268.5. It is notable that the values of two parameters has been selected different, to get a more certain predict result – the diffusion coefficient of the transmission rate of child age group *sβ_c_* is 0.12, and the particle number of sampling to plot the 2D histogram is 1000 in this two cases.

These prediction results suggest that the particle filter model offers the capacity to probabilistically anticipate measles dynamics with a fair degree of accuracy. From the 2D histogram plots, empirical data lying after Prediction Start Time – and thus not considered by the particle filtering machinery – mostly lie within the high-density range of the particles. Notably, in such examples, the particle filter model appears to be able to accurately anticipate a high likelihood of a coming outbreak and non-outbreak. Such an ability could offers substantial value for informing the public health agencies with accurate predictions of the anticipated evolution of measles over coming months.

### Prediction results of classifying outbreak occurrence of the minimal discrepancy model

By incorporating the prediction results of the lowest discrepancy particle filter model *PF_age_15_both_*, we could perform a classification-based prediction of whether the measles will break out or not in the next month. Fig 9 displays the ROC curve showing the prediction results. The Area Under the Curve (AUC) of the ROC curve is 0.89, indicating a favourable classification ability. The confusion matrix at the fixed threshold *θ_k_* = 0.5 is listed in Table 2:

**Fig 9.**
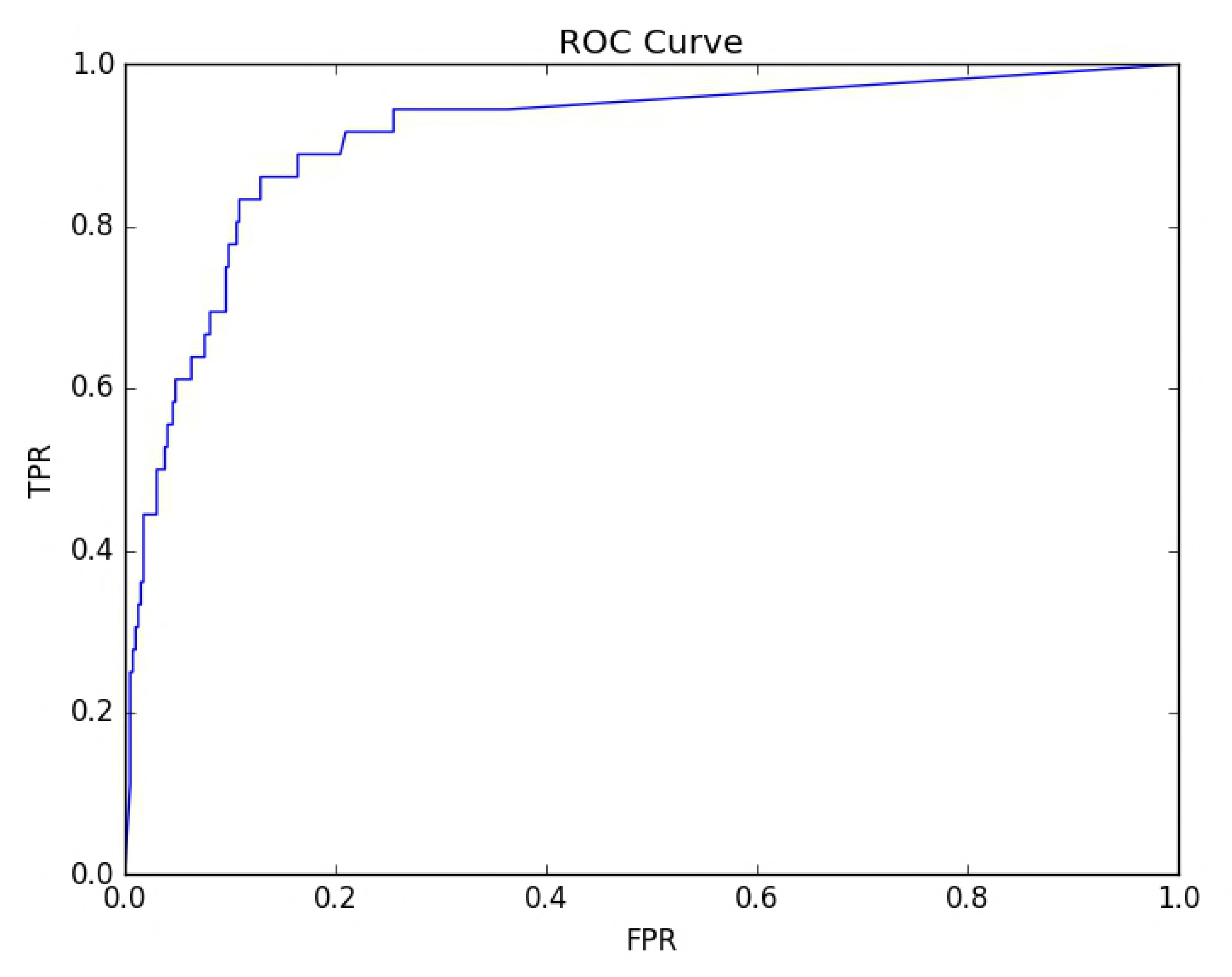
The ROC curve of the prediction classification result of the minimum discrepancy model. The AUC is 0.893.

**Table 2.** The confusion matrix of classifying outbreak occurrence at threshold *θ_k_* = 0.5.

| 432 months | Predicted outbreak | Predicted non-outbreak |
| --- | --- | --- |
| Actual outbreak | 18 | 15 |
| Actual non-outbreak | 18 | 381 |

The threshold $\theta_k = 0.5$ indicates that if there are greater and equal than 50% of particles predict outbreak at each time, then this time is labeled as being in an outbreak status. Otherwise, it is classified as a non-outbreak.

The confusion matrix – Table 2 indicates that across the total timeframe (432 months), there are 33 months that are labeled measles outbreak in empirical dataset. Among these 33 data points, 18 data points are predicted as indicating a non-outbreak. Similarly, among the 399 data points labeled non-outbreak in empirical dataset, 381 data points are predicted as non-outbreak, while 18 data points are predicted as indicating an outbreak.

Fig 10 displays the boxplot of the next month predicted measles counts of sampled particles at the points of the empirical data. The red lines in the boxplot indicate the median of the dataset (the predicted dataset at a specific time). It indicates that the empirical data are located at the range of the predicted data. Thus, we could get the conclusion that the model offers strong predictive performance.

**Fig 10.**
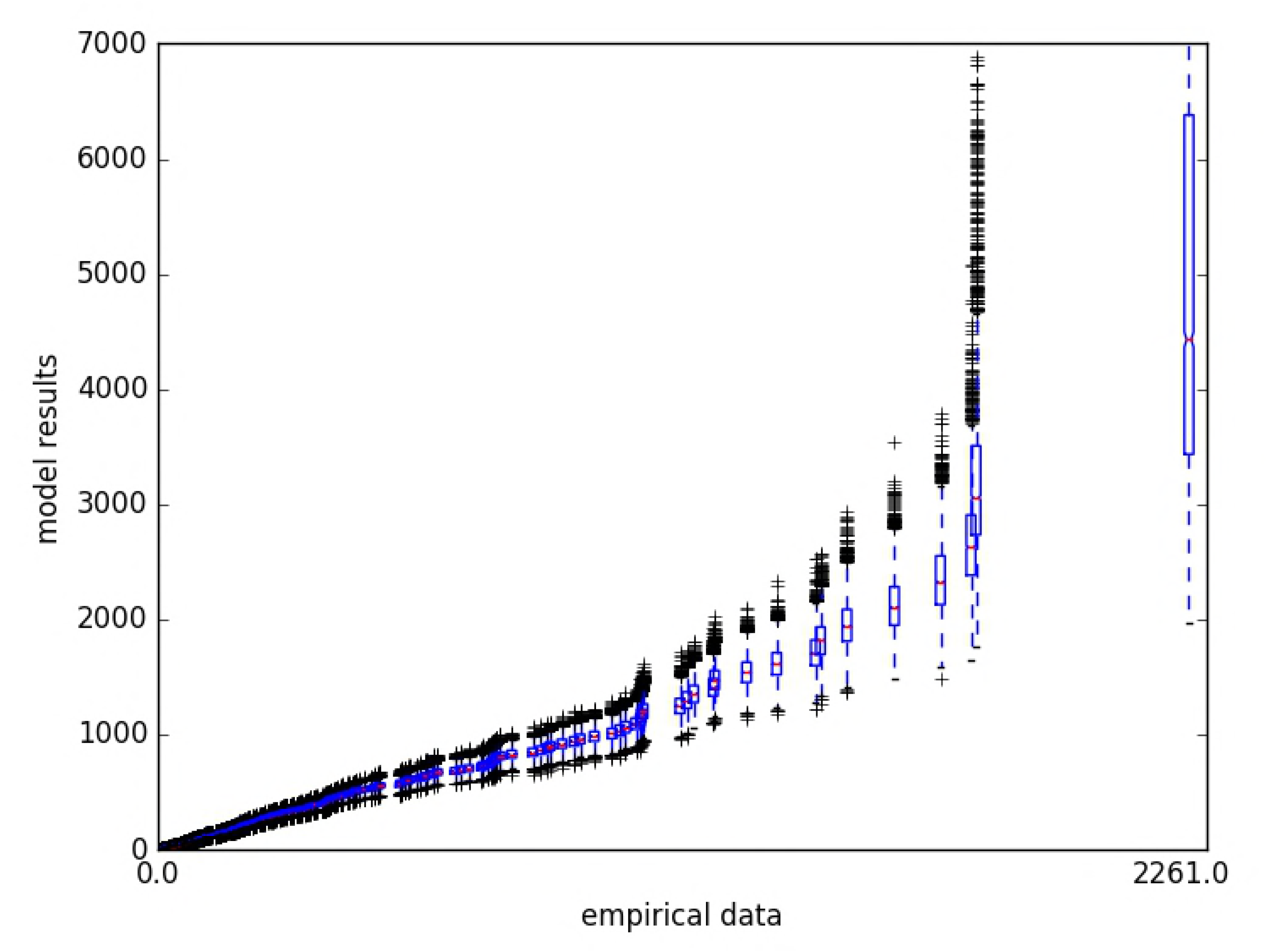
The box plot of the empirical data vs. model calculated results of the minimum discrepancy model.

## Discussion

In this paper we present a new method for tracking the epidemic pattern of measles in low vaccination regions by applying particle filtering with simple measles transmission models, and incorporating noisy monitored data. Particle filtering offers many attractive features for epidemiological models. Firstly, it relaxes the stiff assumptions of normality with respect to the process and observation noise required by older statistical filtering techniques (such as Kalman Filtering); such assumptions are often particularly problematic in epidemiological contexts with small sample counts. Secondly, particle filtering is especially well suited to non-linear models such as that used here, because it foregoes focus on a single Maximum Likelihood Estimate seen in the Kalman Filter – which can be particularly problematic in the context of state uncertainty that can span multiple basins of attraction – and instead samples from a distribution of possible states for a given time-point. In our study, the particle filtering algorithm has mitigated significant weaknesses and simplifications associated with aggregate compartmental models and noisy empirical data. By incorporating ongoing arriving empirical data, the particle filter model has the capability to correct for distortions that accompany compartmental model aggregation, such as assumptions of random mixing and homogeneity. In this dataset, the particle filter offered strong performance in estimating the epidemic pattern of measles and predicting future trends.

Specifically, five particle filter models were investigated in this project. By comparing the results, the strongest predictive performance emerged from the age stratified model whose child age group is defined as including those up to 15 years old, and considering both monthly empirical data of total population and yearly reported cases of each age group. However, testing the models over additional realizations is required in order to validate these results. Finally, we perform prediction analysis based on this best model. The results suggest that particle filtering approaches offer notable strengths in predicting of occurrence of measles outbreak in the subsequent month.

A key benefit of particle filtering lies in its capacity to estimate the latent state of the system – state that cannot be directly measured, but which is jointly implied by the combination of empirical time series and the hypthesized structure of the system, as captured in the mathematical model. It is important to stress that a key motivation for conducting particle filtering to infer latent state in this way lies in the fact that reliable understanding of such latent state is important for estimating the impact of interventions enacted at that point. By estimating the latent state of the system using particle filtering, we can then conduct “what if” scenarios forward from that point, each of which characterize the effects of interventions. Accurate estimation of the state of the system prior to initiation of different intervention strategies will frequently be an important enabler for accurately assessing the differential effects of those interventions.

The particle filtering algorithm in general also has important limitations, such as information loss and particle collapse [39] during the evolutionary process of the particles. These limitations are also inherited in the algorithm applying in this paper that combines particle filtering with a compartmental transmission models. Moreover, the treatment here further suffers from some further challenges. For simplicity, the condensation algorithm is employed in calculating the proposal distribution. However, this algorithm may contribute to a loss of the diversity of particle filtering models. These limitations could be relieved by taking more particles during the calculation, or restricting the particles in appropriate range of changes by selecting the values of parameters in more tightly informed ranges. Finally, a key limitation in terms of practical implications the findings in the developed world reflects the fact that we have focused on prediction in a non-vaccine context; there remains a key uncertainty as to the degree to which the approach proposed here will offer high predictive capacity in the context of sporadic, low attack rate outbreaks characteristic of measles epidemiology in developed countries within the vaccination era.

Much work remains to be undertaken. While particle filtering techniques we investigated in this paper have immediate application in populations with low vaccine coverage (including isolated pockets of population or individuals who refuse vaccination in jurisdictions with otherwise high vaccination coverage), in the future, we will consider vaccination state in the measles particle filter model, to simulate the measles transmission pattern in high vaccination regions in the vaccination era more broadly. Such a model could be helpful for predicting the outbreak of measles in regions suffering from borderline or waning vaccination rates due to vaccine hesitancy, health disparities or other causes. We further plan to apply more powerful techniques, such as Particle Markov Chain Monte Carlo methods that can allow for jointly estimating the latent state of the model and static parameter values whose values are poorly known. Finally, we also plan to investigate more sophisticated means of predicting outbreak occurrence based on particle filtering results. It appears likely that such refinements will further enhance the already strong advantages conferred by particle filtering methods and variants in measles transmission modeling.

We conclude that anticipating the epidemic pattern of measles in low vaccination regions by applying particle filtering with simple measles transmission models so as to recurrently incorporate successive elements of time series of reported case counts is a valuable technique to assist public health authorities in estimating risk and magnitude of measles outbreaks. Such approaches offer particularly strong value proposition for other pathogens with little-known dynamics, critical latent drivers, and in the context of the growing emergence of high-velocity electronic data sources. Additional strong benefits will be realized by extending the application of this technique to highly vaccinated populations.

## Materials and methods

### The introduction of the mathematical SEIR model

This project employs a measles SEIR model [15] as the disease transmission component of the state-equation model in particle filtering. A time unit of months is used, so as to be consistent with the empirical data [29]. Moreover, while the original model lacked age stratification, we also explore an age structured variant of the model. The two model variants used in this paper are introduced as follows.

### Re-dimensionalized aggregate model

This model, which can be found in [15], contains 4 state variables: (S-Susceptible, E-Exposure, I-Infectious, R-Recovered). The original model [15] makes use of a formulation in which each state variable is of unit dimension, representing a fraction of the entire population. However, for the sake of comparison against empirical data, the model in this paper is represented in a re-dimensionalized fashion, with the state variables representing counts of persons. In the first step, we re-dimensionalized the original model. The resulting aggregate compartmental equations are as follows:

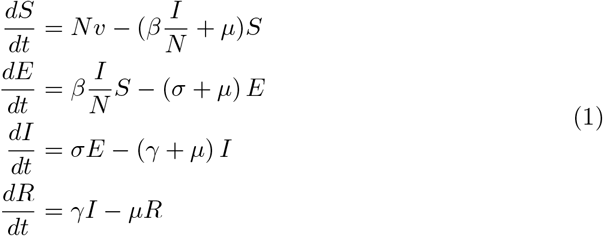

The meaning of the states and parameters are as follows: S, E, I and R are the count of Susceptible, Exposed, Infectious and Recovered people in the population, respectively. N is the total number of people, and N=S+E+I+R. *υ* is the birth rate (of unit 1/Month) and *μ* is the death rate (also of unit 1/Month). *σ*^−1^ and *γ*^−1^ are the mean incubation and infectious periods (in months) of the disease, respectively. *β* is the rate of effective contact between individuals, and reflects both the contact rate and transmission probability (*β* = contact rate × transmission probability), and is thus of unit 1/Month. The population size N was obtained from Saskatchewan during the years from 1921 to 1956. We take the mean of the population across these years to be the value of parameter N. This is associated with the empirical dataset [29]. As noted below, while *β* was treated in [15] as a constant, we took advantage of particle filtering by allowing it to vary over the course of the timeframe explored. The other values of parameters were obtained from [15].

### Re-dimensionalized age structured model

Measles has a severe impact on children’s health and is one of the leading causes of death among young children globally. Measles transmission pattern may be different among different age groups [8]. For example, the age composition of daily contacts may be different for different age groups; children may spend more time with other children and their caregivers, rather than with other adults. Moreover, the rates of contacts sufficiently close to transmitting infection can differ between age groups, such as due to hygienic disparities. To capture such differences, beyond the original model, we created a version of the SEIR model stratified by two age groups: children and adults.

In this variant of the mathematical model, we use subscripts “c” and “a” for a quantity to denote the child- and adult-specific values of that quantity, respectively. We further assume in the demographic model (whose formulation and derivation are introduced in [30]), that the population of each age group (*N_c_,N_a_*) does not change. Similar to the parameter of total population (*N*) in the aggregate model, the mean of the population of each age group across the timeframe in the age pyramid of Saskatchewan [31] is employed as the value of *N_c_* and *N_a_* (where the sum of *N_c_* and *N_a_* equals *N*). The resulting age-structured SEIR model is as follows; readers interested in additional detail are referred to S1 Appendix:

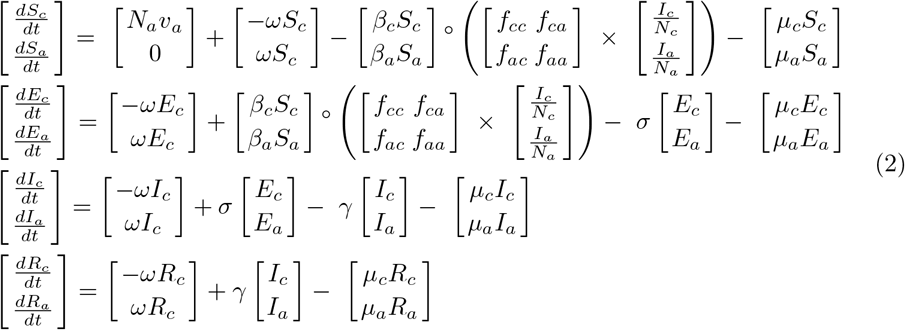

where ° indicates the Hadamard (element-wise) product; × indicates matrix multiplication; 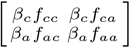 is the contact matrix: *f_cc_* indicates the fraction of children’s infectious contacts that occur with other children; similarly *f_ca_* indicates the fraction of children’s infectious contacts that occur with adults, and *f_ca_*=1−*f_cc_; f_ac_* indicates the fraction of adult’s infectious contacts that occur with children; *f_aa_* = 1 – *f_ac_* indicates the fraction of adult’s infectious contacts that occur with other adults; *ω* is the aging rate out of the age group of children (which carries the same meaning with *c*_1_ in the demographic model in [30]). *υ_a_* is the birth rate for adults, for children, the birth rate is 0. The other parameters hold the same role and values as in the age-aggregated model.

#### The contact matrix model

In the contact matrix 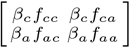, as covered more fully in section “Particle filter implementation in the base model” below, the parameters *β_c_, β_a_* and *f_cc_* are treated as varying across the model time horizon (like the parameter of *β* in Eq (1)). Based upon their values, the other parameters in contact matrix (*f_ca_, f_ac_, f_aa_*) can be calculated as in Eq (3); a detailed mathematical deduction of Eq (3) can be found in S2 Appendix:

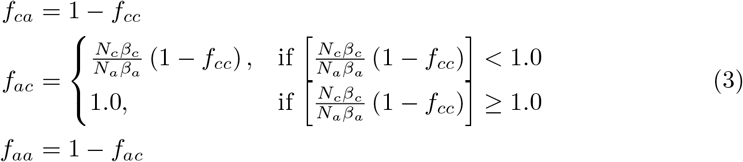

#### The equilibrium demographic model

The population model is listed as follows [30]:

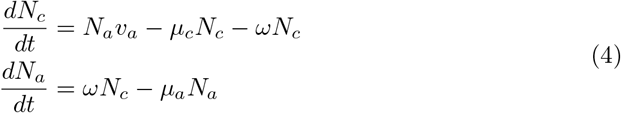

Where *N_c_* is the population of the child age group; *N_a_* is the population of the adult age group; *υ_a_* is the birth rate (applying only to adults); *μ_c_* is the death rate of child age group; *μ_a_* is the death rate of the adult age group.

While measles infection can be lethal, for simplicity, the death rates of all states in the models of this paper are the same; the interested reader is referred to S3 Appendix for additional commentary.

As a result of the assumption of an invariant population size, it follows that death rates for children and adults are as follows; the interested readers are referred to S4 Appendix for a detailed mathematical derivation.

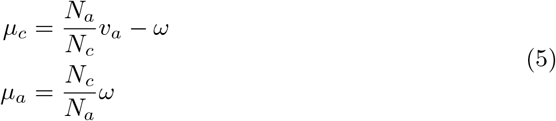

### Introduction to particle filter

Particle filtering is a modern methodology that fits into the broader “statistical filtering” tradition that, as time passes, combines estimates generated by a dynamic model (a model that simulates the evolution of a system over time) with arriving empirical observations. As new data arrives, statistical filtering methods provide a means of arriving at a sort of “consensus estimate” for the current state of the system (as represented by the simulation model) – an estimate that recognizes and balances uncertainty associated with the model (on the one hand) with uncertainty with regards to the observations (on the other). Particle filtering is a very well established contemporary statistical filtering method [32] that is widely employed in fields such as robotics but comparatively new in the sphere of health modeling [16,17,24,26].

Each of the states in the distribution for a given time can be seen as representing a competing hypothesis concerning the underlying state of the system at that time, and the Particle Filtering itself can be viewed as undertaking a “survival of the fittest” of these hypotheses, with fitness determined by the consistency between the expectations of the hypothesis as to what should be observed at observations times, and what is in fact observed at each of those times. Drawing on the theory of Sequential Importance Sampling [16,32,33], this distribution is characterized as a collection of importance-weighted samples, each of which is termed a “particle”. At an intuitive level, a particle’s weight at the current time represents an approximation to the probability of the state represented by that particle in fact obtaining at that time. This weight is, in turn, determined by the consistency of the state being hypothesized by that particle with the observations, as quantified by a likelihood function specifying the likelihood of making a given observation in light of the state captured by the particle.

Interested readers are referred to more detailed treatment in [16,32–34].

### Classifying outbreak occurrence

In this paper, we also investigate the effectiveness of using the particle filtering model in predicting the outbreak of measles in the next time unit (month). The goal of this classification is to map from the reported cases of measles predicted by particle filtering model in the next month to the class of outbreak or non-outbreak. This mapping can be represented by the following equation [32]:

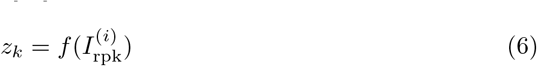

where 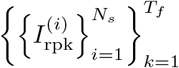 indicates the matrix of the reported cases of measles predicted by the particle filtering model of particle *i* (1 ≤ *i* ≤ *N_s_*) at time *k* (1 ≤ *k* ≤ *T_f_*). *T_f_* is the total running time of the model.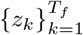 is the vector of the predicted classes – *z_k_* ∈ {0, 1}, where 0 indicates non-outbreak, and 1 indicates outbreak. The value *I*_rpk_ is generated by the particle filtering models. Specifically, the *I*_rpk_ equals *I*_rmk_ in the aggregated particle filtering model (see Eq (11)), and equals *I_rmck_* + *I_rmak_* in the two age groups structured models (see Eq (15)).

Two processes are then used to perform the classification analysis of the results from particle filtering models. In the first process, we define a threshold (*θ*_ik_) of a particle *i* at time *k* above which to consider that particle as positing an outbreak. In the second process, we define a threshold of fraction (*θ_k_*) of particles required to posit an outbreak at time *k* for us to consider there as being an outbreak. Then, the vector of determining whether there is an outbreak of measles in each month – *z_k_* – is calculated.

We further denote 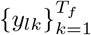 as the empirical vector of whether a measles outbreak indeed obtained at time *k, y_lk_* ∈ {0,1}. The calculation method of *y_lk_* is similar to that of each particle. If the measles reported cases is greater or equal than the threshold *θ*_ik_, the related element in vector *y_lk_* is labeled to be outbreak (the value is 1). Otherwise, non-outbreak (the value is 0).

Finally, to summarize the performance of the classifier, we employ metrics such as the confusion matrix, area under the Receiver Operating Characteristic (ROC) curve, etc. Readers interested in additional detail are referred to S5 Appendix.

### The algorithm of particle filter with next month prediction output

In light of the brief introduction to particle filtering above, the generic particle filter algorithm that we employed in this paper is given as follows [16,33,34]:

1. At time k=0:

1. Sample 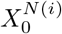 from 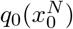;
2. Compute a *weight* for each particle 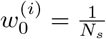. It indicates that the weight at initial time follows the uniform distribution.
2. At time *k* ≥ 1, perform a recursive update as follows:

1. Advance the sampled state by sampling 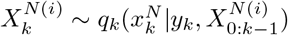 and set 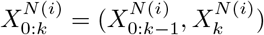;
2. To support classification, output 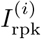 by importance sampling, where 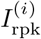 is the sum of all the age groups in the age structured model;
3. Update the weights to reflect the probabilistic and state update models as follows:

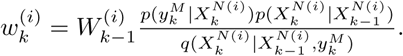 Normalize the weights 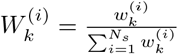
4. Calculate the 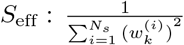
5. If *S*_eff_ < *S_T_* (*S_T_* is the minimum effective sample size – the threshold of resampling), perform resampling to get a new set of 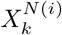. And set the weight of the new particles as 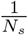.

### Particle filter implementation in the base model

#### The state-space model

The SEIR model is employed as the governing equations underlying the state space model (denoted as *g_k_*) of particle filtering, which is introduced in Eqs (1)–(5). Then each particle at time *k*, noted as 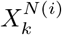, represents a complete copy of the system states at that point of time. Except for the basic states in the SEIR model (S, E, I, R), models of infection transmission are often related to more complex dynamics – such as parameters evolving according to stochastic processes.

#### Aggregate model

In the aggregate model (i.e., the model not stratified by age), three essential stochastic processes are considered. Firstly, we consider changes in the transmissible contact rate linking infectious and susceptible persons, which is represented by the parameter *β*. A second area in which we consider parameter evolution is with respect to the disease reporting process. Specifically, to simulate this process, a parameter – representing the probability that a given measles infectious case is reported *C_r_*, and a state *I_m_* – calculating the accumulative measles infectious cases per unit time (per Month in this paper) – are implemented. Finally, the model includes a stochastic process (specifically, a Poisson process) associated with incidence of infection. This process reflects the small number of cases that occur over each small unit of time (Δ*t*). The stochastics associated with these factors represents a composite of two factors. Firstly, there is expected to both be stochastic variability in the measles infection processes and some evolution in the underlying transmission dynamics in terms of changes in reporting rate, and changes in mixing. Secondly, such stochastic variability allows characterization of uncertainty associated with respect to model dynamics – reflecting the fact that both the observations and the model dynamics share a high degree of fallability. Given an otherwise deterministic simulation model such as that considered here, there is a particular need to incorporate added stochastic variability in parameters and flows capture the uncertainty in simulation results.

To estimate the changing values of these two stochastic parameters (*β* and *C_r_*) and to investigate the capacity of the particle filter to adapt to parameters whose effective values evolve over simulation, the state associated with each particle includes the contact transmission rate *β* and reported rate *C_r_*. Thus, each particle in this project is associated with a state vector *x* = [*S, E, I, R, β, C_r_, I_m_*]^T^.

Reflecting the fact that the transmissible contact rate *β* varies over the entire range of positive real numbers, we treat the natural logarithm of the transmissible contact rate (denoted by *β*) as undergoing a random walk according to a Wiener Process (Brownian Motion) [35, 36]. The stochastic differential equation of transmissible contact rate can thus be described according to Stratonovich notation as:

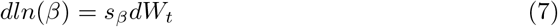

where *dW_t_* is a standard Wiener process following the normal distribution with 0 of mean and unit rate of variance. Then, 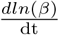 follows the normal distribution with 0 of mean and variance *s_β_*^2^. In this paper, we selected an initial value of *β* following a uniform distribution in the interval [40, 160) across all particles. Unless otherwise noted, the constant value of the diffusion coefficient *s_β_* used is 0.8.

Over the multi-decadal model time horizon, and particularly on account of shifting risk perception, there can be notable evolution in the degree to which infected individuals or their guardians seek care. To capture this evolution, we consider evolution in the fraction of underlying measles cases that are reported (denoted by *C_r_*). Reflective of the fact that *C_r_* varies over the range [0, 1], we characterize the logit of *C_r_* as also undergoing Brownian Motion according to Stratonovich notation as:

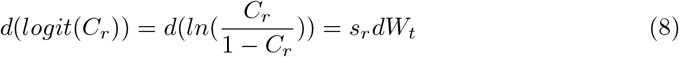

where the initial value of *C_r_* among particles follows a uniform distribution in the interval [0.11, 0.15). The diffusion coefficient (*s_r_*) associated with the perturbations to the logit of *C_r_* over *dt* is selected to be a constant 0.03 across all particles in this paper.

To incorporate the empirical data, we further implement a convenience state *I_m_* with respect to the cumulative count of infectious cases per unit time (Month). The state *I_m_* is different from the state *I*. Specifically, the state of an cumulative count of infectious cases per unit time *I_m_* only considers all the inflows to the infectious state and without all the outflows, to simulate the same process of reporting the measles cases in the real world. The cumulative infectious cases *I_m_* of measles at time *k* is:

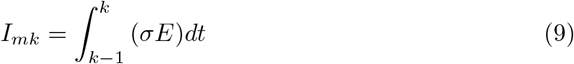

Then, the reported cases at time *k* calculated by the model would be:

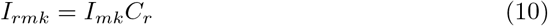

In total, the compartmental model without age stratification is the combination of the classic SEIR model and these three stochastic processes:

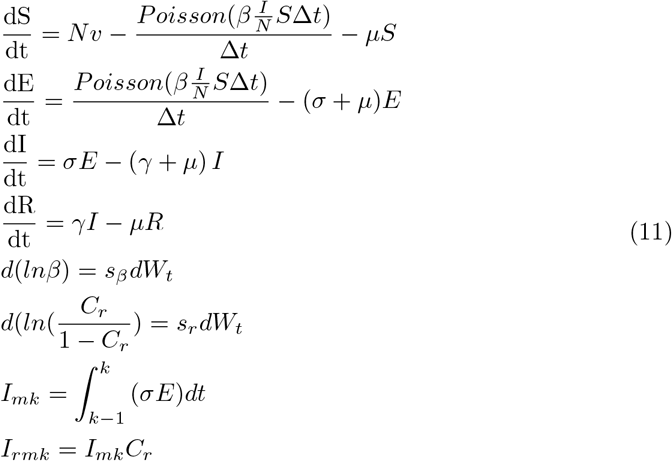

To solve the system above, we made use of Euler integration with a time step of 0.01 Month (i.e., Δ*t*= 0.01 Month in Eq (11)). The same approach was also used in the stratified system presented in Eq (15) below.

#### Age stratified model

In the age stratified model, four stochastic parameters and two extra states are considered dynamically, compared with the original SEIR model (Eqs (2)–(5)). The stochastic process related to incidence of infection is also considered in the age structured state space model, as in the aggregated model above (Eq (11)). The first stochastic parameter is the same as in the aggregate group model: the disease reporting process parameter (*C_r_*), whose dynamics are characterized according to Eq (8). The second is the rate of transmissible contacts between infectious persons and susceptible persons of child age group, which is represented by the parameter (*β_c_*) in the age structured model – Eq (2). The equation and chosen values of (*β_c_*) are the same with Eq (7). The third stochastic parameter (*M_a_*) represents the ratio of the adult age group’s transmissible contact rate (*β_a_*) to that of the child age group (*β_c_*). And we have *β_a_* = *M_a_β_c_*. Reflecting the fact that this parameter represents a non-negative real number, similar to the rate of transmissible contacts (*β*) in the aggregate state space model, we treat the natural logarithm of *M_a_* as undergoing a random walk according to a Brownian motion:

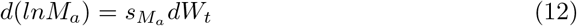

It is a widespread perception that because of limited hygienic awareness and other factors, children are subject to higher rates of transmissible contact than adults. This would suggest that the value of the multiplier *M_a_* normally less than 1.0. Thus, we elected to impose an initial value of *M_a_* across all particles as drawn from a uniform distribution with support [0.2, 1). And, the diffusion coefficient (*s_M_a__*) associated with the evolution of *d*(*lnM_a_*) is chosen to be a constant value of 0.5 among all particles.

The fourth stochastic parameter in the stratified model is the fraction of the contact of children that occurs with other children, denoted as *f_cc_*. This parameter appears in the contact matrix, and varies over the range from 0 to 1. As a result, the dynamic process for *f_cc_* is similar to the disease report rating *C_r_* with the Eq (8), specifically:

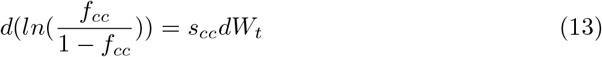

The initial value of *f_cc_* we employed follows a uniform distribution in the interval [0.2, 1.0). We assumed a constant value of 0.2 as the diffusion coefficient (*s_cc_*) associated with the logit of *f_cc_*.

Finally, the new state of cumulative infectious count per unit time is implemented similar to aggregate model (Eq (9)), except for its division into two distinct states according to stratification into two age groups (*I_mc_* and *I_ma_*). The discrete time equations of *I_mc_* and *I_ma_* and reported infectious count per unit time in model *I_rmc_* and *I_rma_* at time k are as follows:

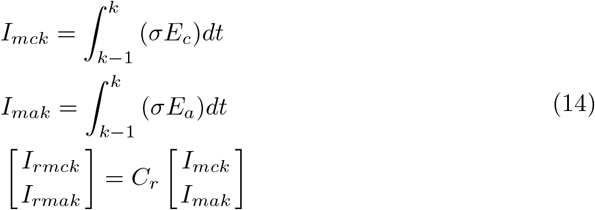

The state vector *x^N^* is [*S_c_, S_a_, E_c_, E_a_, I_c_, I_a_, R_c_, R_a_, β_c_, M_a_, f_cc_, C_r_, I_mc_, I_ma_*] in the age stratified model, and N equals 14. The complete set of state equation for the age-stratified model is given in (15):

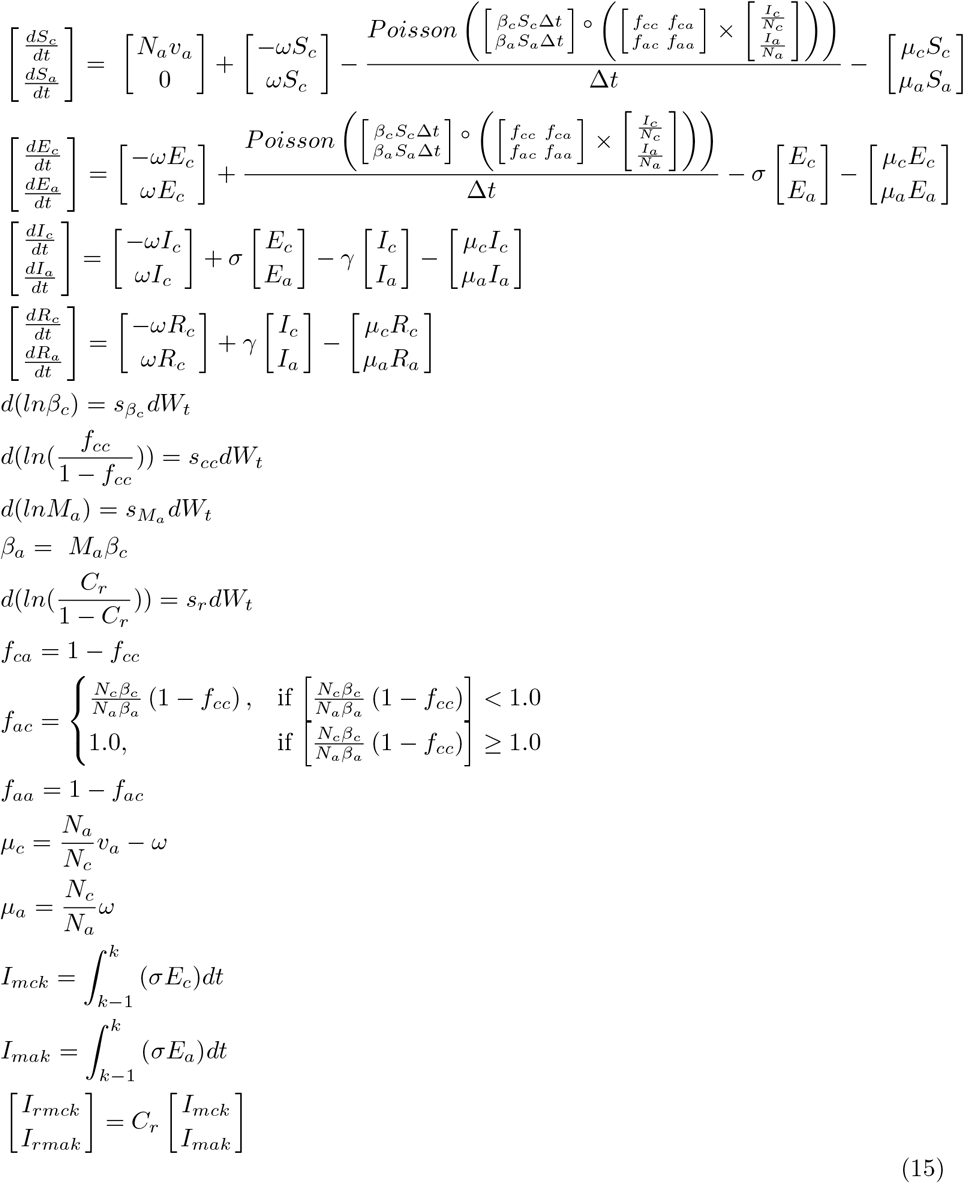

Reflective of the structure of the age group stratification in available data, it is notable that we have considered two different age group configurations in this paper: one where the child age group includes those up to 5 years old (*M_age_5_*) and another where it includes those up to 15 years old (*M_age_15_*).

#### The measurement model

As introduced in particle filtering tutorials [33, 34], the measurement model characterizes the relationship between the measured data and the model. In this paper, we denote the measurement vector as 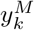, where *M* indicates the length of the measurement vector.

#### Aggregate model

In the aggregate model, the measurement vector 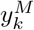 is one-dimensional (representing empirical dataset of monthly reported measles infected cases), that is, M equals 1. We denote the value of empirically reported cases as *I_em_*. Then, at time *k*, the measurement model can be represented as:

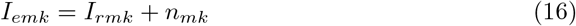

where *I_rmk_* is calculated by the state space model of Eq (11), and *n_mk_* is the measurement noise related to the monthly reported cases.

#### Age stratified model

In the age stratified model, empirical observations can include three components. Thus, in the measurement vector 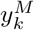, M is 3 in age stratified model. In addition to the empirical data associated with monthly reported cases, empirical data further include annual reported cases for each of the two age groups (children and adults). The measurement model of age stratified model in this paper can thus be represented as:

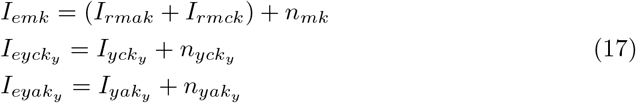

where *I_emk_* is the same as in the aggregate model in Eq (16); *I_rmak_* and *I_rmck_* are calculated in the state space model of the age stratified model in Eq (15); *I_eyck_* consists of the annual measured cases of child age group, while *I_eyak_y__* represents the annual measured cases of adult age group; *I_yck_y__* and *I_yak_y__* are the annual reported cases calculated by the state space model of Eq (15) of children and adult age group, respectively; *n_yck_y__* and *n_yak_y__* are the measurement noise associated with these two age groups.

It is notable that the subscript *k_y_* indicates annual time points, while the unit of time in the models in this paper is month. Thus *I_yck_y__* and *I_yak_y__* could be obtained by the sum of *I_rmak_* and *I_rmck_* in the model each year.

### The proposal distribution

The Condensation Algorithm [32, 37] is applied in this project to implement the particle filter model. It is the simplest and most widely used proposal distribution, and consists of a proposal distribution that is the same as the prior [32, 33].

### Likelihood function

In this project, the observed data is of two types – the monthly reported incidence case count of measles and annual reported cases within different age groups. As previously introduced, the measured data is the reported cases of measles in this paper. The likelihood function 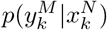 describes such a reporting process, and specifies the probability that a given measles case in the dynamic model will be measured. We followed several past contributions [16,17,20,38] in selecting the negative binomial distribution as the basis for the likelihood function, which allows for greater robustness than the classic binomial distribution. Readers interested in additional detail are referred to S6 Appendix.

The equation associated with the likelihood function is thus as follows:

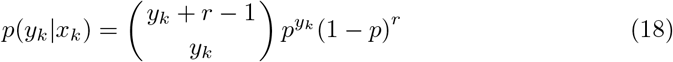

where *y_k_* is the empirical data (reported measles cases) at time 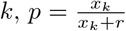 representing the probability that a given measured reported case is in fact a true reported incident case, and *x_k_* is the (integer rounded) incident cases resulting from the dynamic model at time *k*. *r* is the dispersion parameter associated with the negative binomial distribution. In all scenarios reported in this paper, the value of *r* is chosen to be 10.

### Aggregate model

In the aggregate model, because the model lacks the capacity to distinguish between individuals within different age groups as necessary to compare to the yearly age-stratified reported values, the measured data is a one-dimensional vector consisting of the monthly reported cases. It indicates that the weight update rule (likelihood function) of the aggregate model could simply achieved by calculating the value of *p*(*y_mk_*|*I_rmk_*), where (*y_mk_* is the empirical data as given by the monthly reported measles cases at time *k*, and *I_rmk_* is the reported cases calculated by the dynamic model.

### Age stratified model

In the age stratified model, the weight update rule is similar to that in the aggregate model, except for the update associated with the close of each year. Specifically, the weights of particles associated with the age stratified model from January to November are only updated by the monthly empirical data – monthly measles reported cases at each time (using the likelihood function given in Eq (18)). However, the weight at the end of the last month (December) of each year is updated by the combination of three parts. The likelihood formulation of age stratified model is listed as follows:

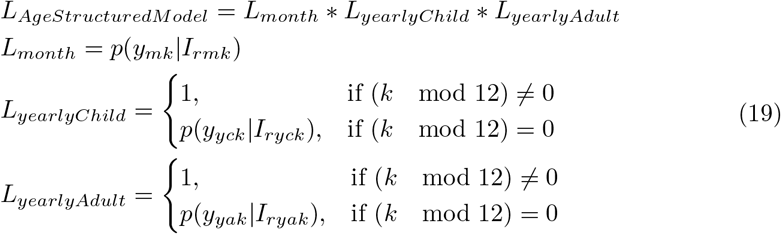

where *L_month_* is the likelihood function based on the monthly empirical data for the total population. The other two likelihood functions reflect the fact that annual totals are available on an age-specific basis at year end. *L_yearlyChiid_* is the likelihood function based on the yearly empirical data for the child age group. *L_yearlyAdult_* is the likelihood function based on the yearly empirical data of the adult age group.

## Evaluating particle filter performance

To assess the accuracy of particle filtering for robust estimation of model states, it is essential to evaluate the estimation and predictive capacity of the particle filtered models. In this project, we therefore sought to calculate the discrepancy at each observation time (Month in this paper) [16] between the model generated data and empirical data, using a linear measure. Reflecting the dual sources of data employed, the discrepancy includes monthly discrepancy and yearly discrepancy, with the total discrepancy representing the sum of these quantities.

Typically, there will be thousands of particles included in each model run. To calculate the discrepancy of particle filter results by incorporating empirical data across all time points, we sample n particles by importance sampling for each such time. The monthly discrepancy of each time is simply the Root Mean Squared error (RMSE) between the monthly empirical data at that time and the related data calculated by the particle filtering model [16, 22]. To get the yearly discrepancy of each time (here, successive Months), the RMSE was calculated for each age group of each year (similary to monthly discrepancy). Then the yearly discrepancy is the sum across all age groups of the yearly RMSE over 12 (to convert the unit to Month).

Moreover, to investigate the predictive capability and efficiency of particle filter model, we defined a variable “Prediction Start Time”, denoted by *T*\*; it indicates the time 1 ≤ *k* < *T*\* up to which the weights of particles are updated based on the observed data. When *k* ≥ *T*\*, the particle filtering ceases – the weights of particles are no longer updated and no further re-sampling occurs. During that period (i.e., following the Prediction Start Time), particles simply continue to evolve according to the state space model shown in Eq (11) or Eq (15) (depending on whether an age-specific model is being applied). For such a Prediction Start Time *T*\*, the model calculated a prediction discrepancy using a simple variant of the strategy of the discrepancy used in considering all the time frame, but limited to considering only times *T*\* and larger.

## Empirical dataset

### Measles reported cases

In this project, the empirical data consists of the time series from 1921 to 1956 of reported measles cases for the mid-western Canadian province of Saskatchewan. These aggregate data are obtained from Annual Report of the Saskatchewan Department of Public Health [29]. We have employed two datasets: monthly reported measles cases aggregated across the total population and yearly reported cases in each of different age groups. In the empirical yearly dataset, these yearly reported cases are split into different age groups. In a small minority of years (from 1926 to 1941), the age categories present in the reported data do not correspond neatly to the age group categories in the models (considering children as being those within their first 5 years or first 15 years). For these cases, we split them into the age categories of the models proportionally. Readers interested in additional detail are referred to S7 Appendix.

We consider the pre-vaccination era of measles within Saskatchewan to study the natural dynamics of measles in low vaccination areas. The time of the monthly empirical data is from Jan. 1921 to Dec. 1956, with the monthly dataset offering a total of 432 records. The yearly age specific data are from 1925 to 1956, reflecting the fact that reporting of age specific data is only started in 1925. Every record contains three features – date, measles reported cases and population size. To make them consistent with the total population size of the dynamic model (863,545), the empirical data are normalized to the same population size of the model. The normalized monthly empirical data are shown in Fig 13; it can be readily appreciated that the time series demonstrates the classic patterns of waxing and waning incorporating both stochastics and regularities characteristic of many childhood infectious diseases in the pre-vaccination era.

**Fig 11.**
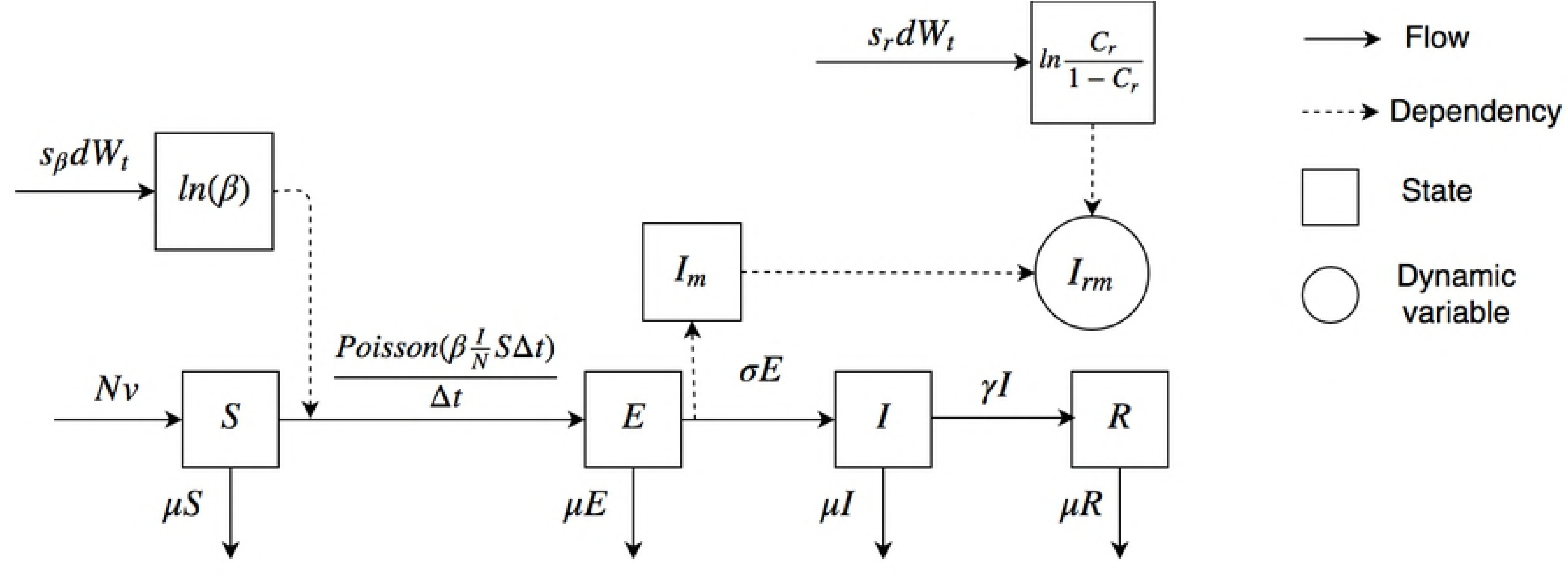
The mathematical structure of the particle filtering aggregate model.

**Fig 12.**
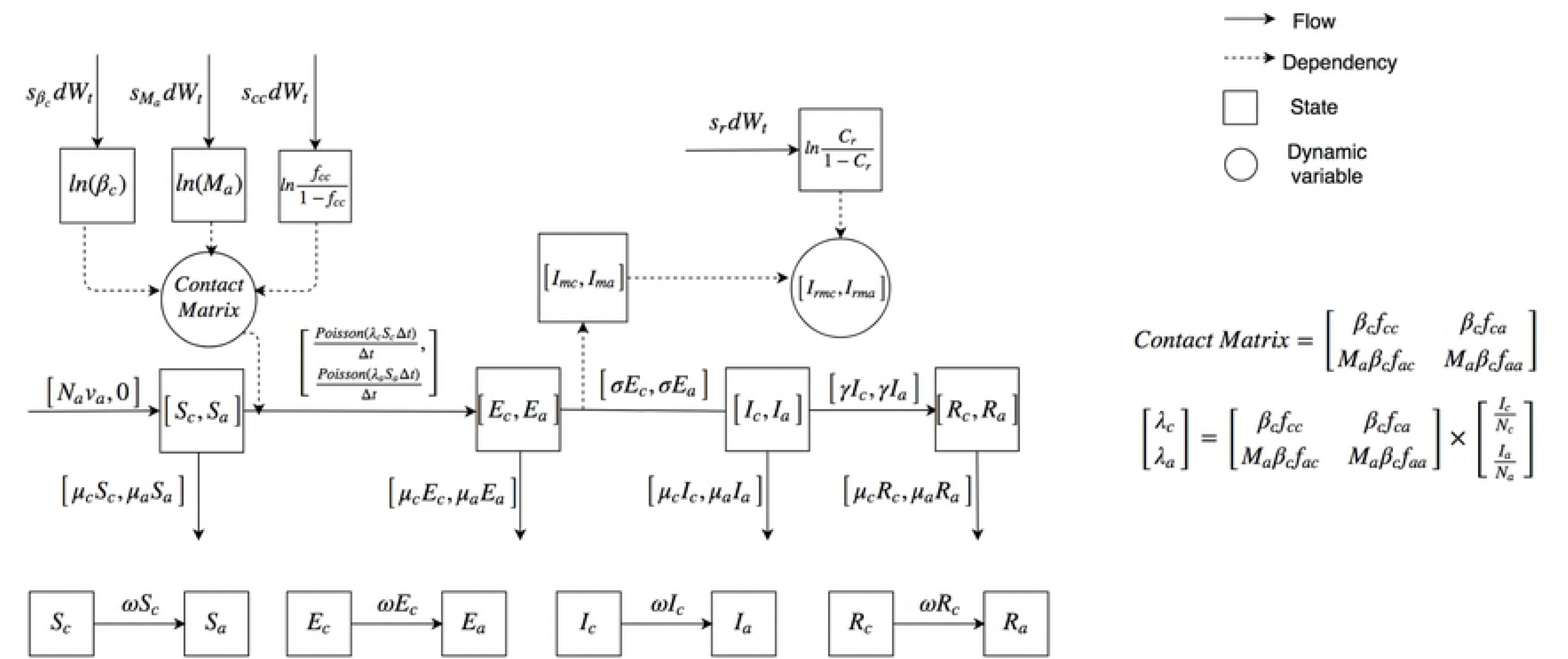
The mathematical structure of the particle filtering age stratified model.

**Fig 13.**
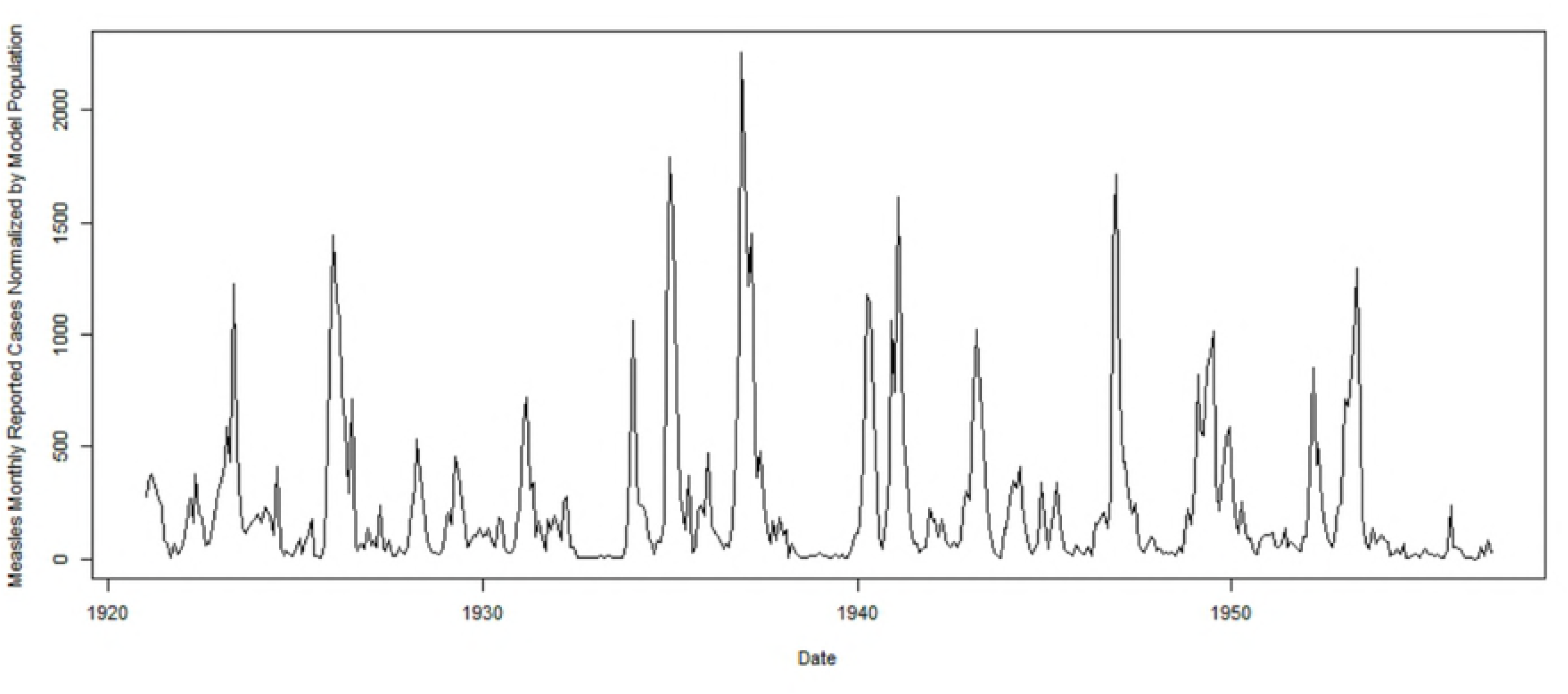
The monthly reported measles cases in Saskatchewan from 1921 to 1956. The values given are normalized by the population employed in the model (863,545).

### Population introduction

In this project, the parameters of the population play a significant role in the models (Eqs (1)–(5)). The parameters (*N, N_c_, N_a_*) related to the population are abstracted from the empirical population data of Saskatchewan from 1921 to 1956 [31]. From 1921 to 1956, the empirical population lies in the interval from 757,000 to 932,000. And the empirical total population of Saskatchewan in these years does not exhibit drastic fluctuation [31]. Thus, we let the model population remain constant at 863,545, which represents the mean population over that interval. Also, the monthly total and yearly age groups’ empirical measles reported cases are normalized to the model population. It is notable that we employ an equilibrium population model – the total population and population among each age group will stay the same, rather than change. Similarly, the values of the population in each age group also employs the average population among 1921 to 1956 in their age group as given by the age pyramid [31].

### Parameters

The important fixed parameters in the models are *γ*^−1^, *σ*^−1^, *υ, μ, v_a_, N, N*_*c*15_, *N*_*a*15_, *N*_*c*5_, *N*_*a*5_. The values of birth rate were also obtained from the Annual Report of the Saskatchewan Department of Public Health [29]. The values of parameters of *γ*^−1^ and *σ*^−1^ are as given by [15]. Moreover, to compare the results, we have built two types of age structured models – one where the lower age group consists of children below 5 years of age (with population in child age group denoted as *N*_*c*5_, and population of adult age group denoted as *N*_*a*5_), and another in which children consist of individuals below 15 years of age (population of age groups denoted as *N*_*c*15_ and *N*_*a*15_, respectively). Thus, the birth rates are different among these two types of models (denoted as *υ*_*a*5_ and *υ*_*a*15_ respectively), to let all the models have a similar birth population per unit time. Finally, all the compartmental parameters are specified at Table 3. The initial value of all stocks in the particle filtering models are given in S1 Table.

**Table 3.** Table showing the value of parameters.

| Parameter | Value | Units |
| --- | --- | --- |
| $\gamma^{-1}$ | 5 | Day |
| $v$ | 0.03 | 1/Year |
| $\mu$ | 0.03 | 1/Year |
| $N$ | 863,545 | Person |
| $\sigma^{-1}$ | 8 | Day |
| $v_{a15}$ | 0.045 | 1/Year |
| $v_{a5}$ | 0.034 | 1/Year |
| $N_{c5}$ | 98,743 | Person |
| $N_{a5}$ | 764,802 | Person |
| $N_{c15}$ | 286,537 | Person |
| $N_{a15}$ | 577,008 | Person |

## Supporting information

S1 Appendix. The mathematical deduction of the age structured epidemiology model.

S2 Appendix. The mathematical deduction of the contact matrix model.

S3 Appendix. The complementary comments of the parameter of death rate.

S4 Appendix. The mathematical deduction of the death rate.

S5 Appendix. The further introduction of the classification method.

S6 Appendix. The shortcoming of choosing the binomial distribution as the likelihood function.

S7 Appendix. The further introduction of split the measles yearly reported cases to each age group.

S1 Table. The Initial Values of All Particle Filtering Models.

## Acknowledgments

We thank Weicheng Qian and Anahita Safarishahrbijari for the suggestions of building models; we also thank Lujie Duan for the help of debugging the models.

